# Cardiolipin Remodeling as a Regulatory Switch in Osteogenesis: Insights from Multimodal Metabolic Imaging and Molecular Profiling

**DOI:** 10.64898/2025.12.22.696122

**Authors:** Maria Iftesum, Praveen Kumar Guttula, Kirti Agrawal, Subhrajyoti S Kundu, Sreyashi Das, Fabrizio Donnarumma, Ram Devireddy, Manas Ranjan Gartia

**Affiliations:** Department of Mechanical and Industrial Engineering, Louisiana State University, Baton Rouge, LA 70803, USA; Department of Chemistry, University of Florida, Gainesville, FL 32611, USA; Mass Spectrometry Facility, Department of Chemistry, Louisiana State University, Baton Rouge, LA 70803, USA

## Abstract

Cardiolipin (CL), a mitochondria-specific phospholipid, plays a fundamental role in respiratory chain organization and bioenergetic efficiency, yet its contribution to osteogenic differentiation is poorly defined. Here, we used a multimodal approach integrating untargeted LC–MS lipidomics, Raman spectroscopy, fluorescence lifetime imaging microscopy (FLIM), and structural imaging to investigate CL remodeling during human adipose-derived stem cell differentiation. Lipidomics revealed a selective enrichment of highly unsaturated CL species, accompanied by transcriptional upregulation of the cardiolipin biosynthetic and remodeling enzymes CDS1/2, PGS1, CRLS1, TAZ, and HADHA. Lipidomics also revealed a time-dependent increase in membrane-associated lipids including phosphatidylcholine (PC), serine, and phosphatidylinositol (PI). These lipids were implicated in supporting mitochondrial membrane expansion, oxidative phosphorylation, and signaling processes critical for osteoblast maturation. Spatial imaging techniques confirmed cardiolipin accumulation and redistribution in differentiated cells, while Raman-based direct classical least squares (DCLS) analysis provided label-free mapping of lipid species. Gene Ontology (GO) enrichment and protein–protein interaction network analysis further identified biological pathways related to bone remodeling, cardiolipin metabolism, and osteoblast-specific signaling. Fluorescence Lifetime Imaging Microscopy (FLIM) data revealed a metabolic shift from glycolysis to oxidative phosphorylation during differentiation, supported by structural and gene expression evidence. These changes temporally coincided with matrix mineralization and collagen organization, linking CL metabolism to both cellular bioenergetics and extracellular matrix production. Our findings identify cardiolipin remodeling as a metabolic checkpoint in osteogenesis and suggest that targeted modulation of CL pathways may provide new therapeutic strategies for enhancing bone regeneration.

## INTRODUCTION

Osteogenic differentiation is a finely tuned biological process that results in the conversion of mesenchymal stem cells (MSCs) into osteoblasts that produce bone through the deposition and calcification of extracellular matrix (ECM) [1, 2]. This process is composed of separate stages that are governed by multiple interconnected pathways of signaling, transcription factors, and metabolic changes [3, 4]. Human adipose-derived stem cells (hASCs) are uniquely advantageous for regenerative medicine and bone tissue engineering purposes due to their accessibility, ease of procurement, and robust osteogenic capacity [5, 6], making them valuable for treating conditions such as osteoporosis and bone defects [7, 8].

Most of the research has been conducted on the genetic and proteomic levels, with emphasis on transcription factors such as Runt-related transcription factor 2 such as (RUNX2), Osterix (OSX), and activating transcription factor 4 (ATF4) as well as their downstream markers alkaline phosphatase (ALP), osteopontin (OPN), and osteocalcin (OCN) [9, 10]. However, more recent studies show that lipid metabolism is a vital yet understudied process that affects osteogenic differentiation through energy metabolism, signaling pathways, and structural elements [11–13]. Lipid metabolism plays a vital role in osteogenesis by supplying energy, supporting cellular structure, regulating signaling pathways, and maintaining overall bone health [14]. During osteoblast differentiation and bone formation, fatty acid oxidation becomes a key energy source, complementing or even surpassing glucose metabolism, especially under high-demand conditions such as growth or fracture repair [15]. Lipids such as cholesterol and phospholipids are essential components of cell membranes, ensuring structural integrity and facilitating interactions within the bone microenvironment [16, 17]. Additionally, lipid-derived signaling molecules influence critical pathways like Wnt/β-catenin and PPARγ, which regulate osteoblast differentiation and the balance between osteogenesis and adipogenesis [15, 18]. Disruptions in lipid metabolism, such as excessive lipid accumulation or dyslipidemia, can impair bone remodeling and are associated with decreased bone mineral density and increased fracture risk [19]. Despite its acknowledged relevance, the detailed analysis of lipid metabolic alterations in osteoblast differentiation is scarce and calls for further examination [20–22].

The development of osteogenic cells depends significantly on lipid metabolism as a critical mechanism that affects both energy generation and cell communication as well as tissue stability. Lipids are essential for various cellular functions such as membrane structure and function, signal transduction and metabolic adjustments that support osteoblast function and bone matrix mineralization [13, 23]. Specific lipid classes, such as phospholipids, enhance cellular membrane modification and enhance signal transmission, whereas sphingolipids are involved in the regulation of signaling pathways that control osteoblast survival and differentiation [24].

Current research has highlighted cardiolipin as a specialized mitochondrial phospholipid with essential functions in mitochondria, and cardiolipin is an essential phospholipid for mitochondrial function and structure [25–27]. Cardiolipin is essential for the efficient function of oxidative phosphorylation (OXPHOS) and production of ATP which are needed to supply the high energy needs of osteoblast maturation [11, 28]. Changes in cardiolipin content directly impact the bioenergetic properties of mitochondria, impacting energy metabolism and cellular health and thus making it a potential biomarker for monitoring the stages of osteoblast differentiation [29]. However, chemical mapping of these lipids during osteogenic differentiation process has not been performed before.

In this study, we combined Raman spectroscopy with LC-MS lipidomics and FLIM and SHG and HSI and gene expression analysis to demonstrate that osteogenic differentiation depends on cardiolipin and phosphatidylcholine enriched lipid metabolic reprogramming. We also examine the metabolic shift during the differentiation process. Our approach would help reveal how lipid metabolism regulates osteoblast function. These findings might lead to new treatments to improve bone formation in metabolic bone disorders.

## RESULTS

To investigate lipid remodeling during osteogenic differentiation, we profiled global lipid classes and cardiolipin (CL) species composition. Fig. 1a shows that the variation of lipids fold changes of control vs. differentiated cells in week 1, week 2 and week 3. For each week the values are averaged over n=2 replicates. Heatmap analysis of major lipid classes revealed distinct abundance patterns that separated control from differentiated cells (Fig. 1a). At the species level, Cardiolipins showed extensive remodeling, with several molecular species significantly altered during differentiation (Fig. 1b).

**Fig. 1.**
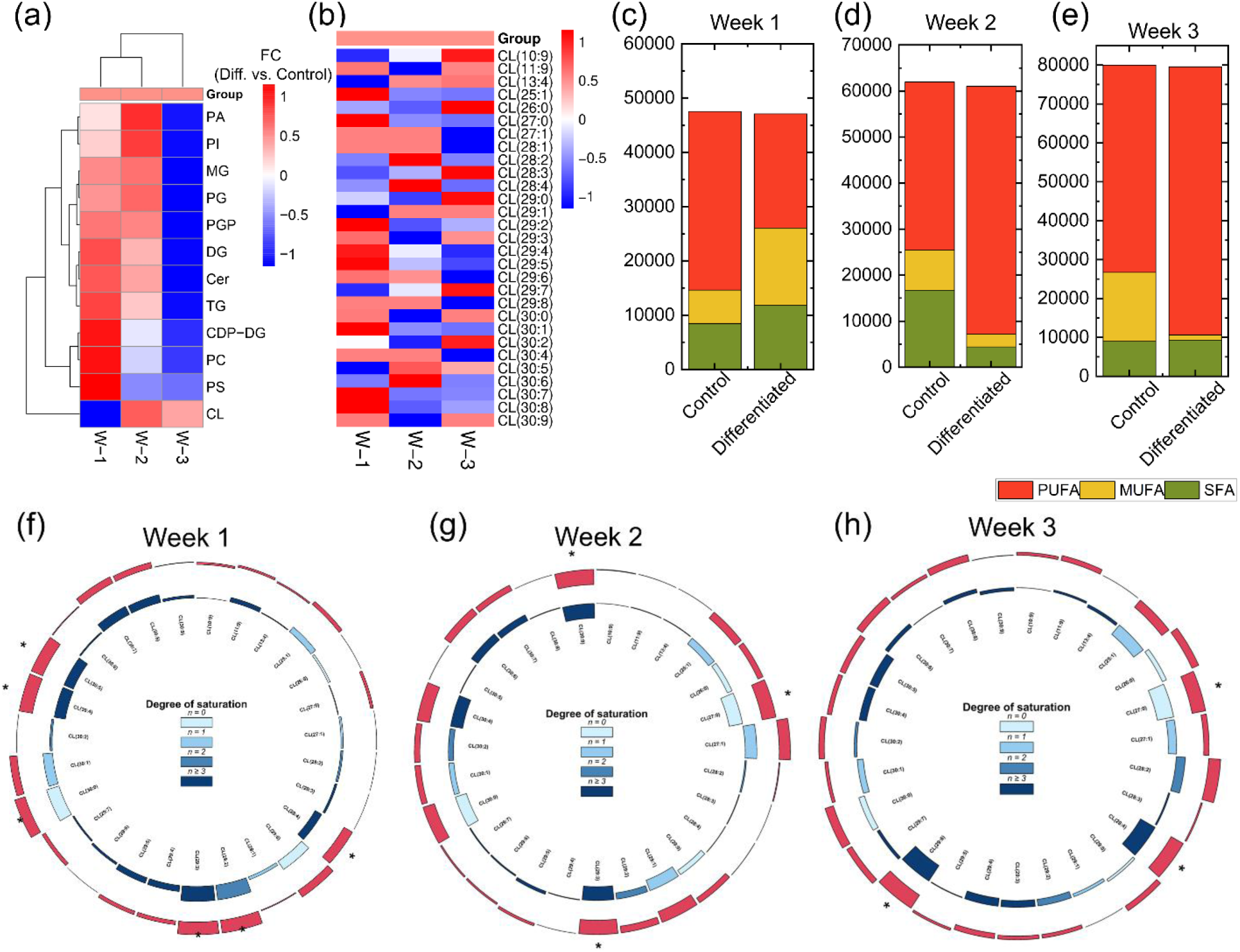
Comprehensive lipidomic analysis highlights remodeling of cardiolipin and fatty acid composition during osteogenic differentiation: (a) Heatmap showing the Fold change (Differentiated vs. Control) patterns of major lipid classes for week 1 (W-1), week 2 (W-2) and week 3 (W-3) of cell culture; (b) heatmap of individual cardiolipin (CL) species fold change (FC) (Diff vs. Control) illustrating distinct remodeling during differentiation. For each week the values are averaged over n=2 replicates; (c–e) stacked bar plots showing changes in the distribution of polyunsaturated fatty acids (PUFA), monounsaturated fatty acids (MUFA), and saturated fatty acids (SFA) in control and differentiated cells for week 1, week 2 and week 3 of cell culture; (f–h) circos plots depicting the degree of fatty acyl chain unsaturation in cardiolipin species, highlighting enrichment of highly unsaturated species and depletion of saturated species during differentiation, with asterisks marking significantly altered species (p < 0.05). The inner ring depicts the fatty acyl elongation index for each lipid, with bar colors reflecting unsaturation level: light blue for saturated (n = 0), sky blue for monounsaturated (n = 1), steel blue for di-unsaturated (n = 2), and navy for polyunsaturated chains (n ≥ 3). In the outer ring, log2-fold changes of median-normalized lipid levels are shown, where upward red bars indicate increased abundance in differentiated cells and inward bars mark reductions.

Examination of fatty acid composition demonstrated a redistribution among polyunsaturated (PUFA), monounsaturated (MUFA), and saturated fatty acids (SFA). Differentiated cells exhibit a relative increase (∼1.5x) in PUFA-containing lipids (Fig. 1(c–e)) from week 1 to week 3. The PUFA in control was higher compared to the differentiated cells in week 1, while it was reversed for week 2, week 3, with PUFA of differentiated cells higher compared to control ones. The MUFA of differentiated cells was higher in week 1 compared to control while they were lower in week 2 and week 3. The SFA also showed a different distribution from week 1 to week 3 of culture. SFA was higher in week 1 differentiated cells compared to control in week 2, but remain similar in week 3.

Shown in Fig. 1f–h are circos plots of cardiolipins (CLs) across three weeks of osteogenic differentiation. CLs are annotated by “lipid subclass” subsequently by the “total fatty acyl chain length:total number of unsaturated bonds”. The species in each CL class are ordered from fully saturated to highly polyunsaturated and, within each subgroup, from shortest to long. The outer ring shows changes in abundance (log2FC) of each lipid, To most optimally visualize altered acyl chain length, we expressed the abundance of CL species in differentiated and control relative to the shortest CL species of each saturation subclass (the inner ring shows the elongation index (EI)), which reflects whether fatty acyl chains are getting longer (positive values) or shorter (negative values). In Week 1, many CLs increased, especially the longer-chain, polyunsaturated ones such as CL(29:2), CL(29:3), and CL(30:0). The EI confirmed strong elongation in these species, although some very unsaturated CLs (like CL 30:4 and CL 30:5) were shortened. In Week 2, the pattern was more mixed. Some CLs (e.g., CL 25:1, CL 27:0, CL 29:3) decreased, while others (like CL 29:5, CL 30:6, and CL 30:7) increased. EI values also showed this balance: certain species were elongated (CL 27:1, CL 30:6, CL 30:7), while others were shortened (CL 27:0, CL 29:3, CL 30:9). By Week 3, several highly polyunsaturated CLs (CL 25:1, CL 27:0, CL 28:4, CL 29:6) were strongly reduced, while a few species (CL 11:9, CL 28:2, CL 30:0, CL 30:2) remained higher. EI values showed less elongation overall, with many long PUFA cardiolipins being shortened. Overall, the data reveal a three-phase remodeling process:

Week 1 – expansion of long-chain, PUFA cardiolipins through elongation.

Week 2 – a mixed phase with both elongation and shortening.

Week 3 – reduction of elongated PUFA cardiolipins, leaving a more stable pool of shorter or moderately unsaturated species.

This pattern suggests that mitochondria first build up a diverse cardiolipin pool to fuel early differentiation and later shift toward protecting membranes from oxidative damage during mineralization.

To dissect pathway-specific lipid remodeling during osteogenic differentiation, we analyzed phospholipid intermediates along the PA–CDP-DG–PGP–cardiolipin biosynthetic axis. In PA (Fig. 2a, d), total abundance increased by roughly ∼2× in differentiated cells, driven by rises in MUFA (∼2–2.5×) and PUFA (∼1.5–2×), while SFA showed only a modest change (∼+10–20%). The total CDP-DG and PG pool decreased signifying conversion to CL. In CDP-DG (Fig. 2b, e), the total pool declined ∼10–20%, with MUFA decreasing ∼30–40% and PUFA decreasing ∼10–20%; SFA remained approximately stable (0–10% change). In PGP (Fig. 2c, f), the total pool fell ∼10–20%, with a pronounced reduction in MUFA (∼60–70%) and a more moderate drop in PUFA (∼15–25%), whereas SFA changed little (∼0–10%). Consistent with increased channeling of PA toward cardiolipin, transcripts for CDS1/2, PGS1, PGPP1/2/3, and CRLS1 were elevated (Fig. h), and pathway modeling (Fig. i) showed positive Z-scores (DG→PA, Z = 2.666) along reactions funneling PA→CDP-DG→PGP→CL with suppression of competing routes. Collectively, the data indicates that osteogenic differentiation expands a PUFA- and MUFA-rich PA reservoir, while CDP-DG and PGP pools contract, consistent with enhanced flux into cardiolipin biosynthesis.

**Fig. 2.**
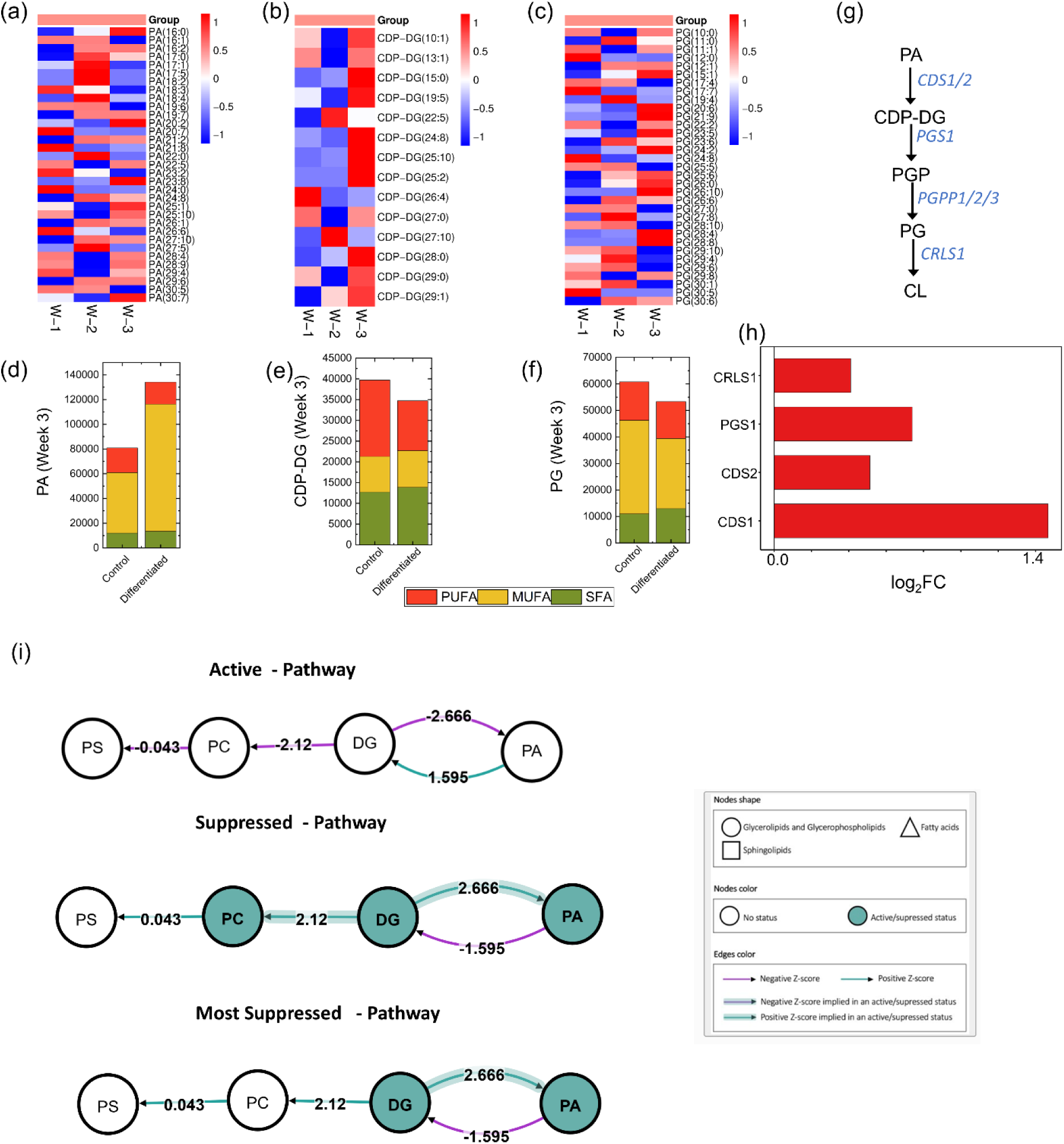
Pathway-specific remodeling of phospholipid metabolism during osteogenic differentiation: (a–c) Heatmaps of (a) phosphatidic acid (PA), (b) CDP-diacylglycerol (CDP-DG), and (c) phosphatidylglycerophosphate (PGP) species using FC (Differentiated vs Control) showing distinct shifts between control and differentiated cells among week 1, week 2 and week 3; (d-f) corresponding stacked bar plots illustrate the distribution of total polyunsaturated (PUFA), monounsaturated (MUFA), and saturated fatty acids (SFA) in week 3 between control and differentiated cells for (d) PA, (e) CDP-DG and (f) PG; (g) Biosynthesis pathway of CL from PA with the corresponding enzymes involved; (h) Expression profile of key enzymes regulating the cardiolipin biosynthesis pathway using RNAseq data of week 3 Differentiated vs. Control osteogenic stem cells; () (CDS1/2, PGS1, PGPP1/2/3, CRLS1). (i) Biochemical pathway modeling of lipid interconversions highlights activation and suppression of specific reactions, with active nodes and positive Z-scores (teal) denoting enhanced flux through PA biosynthesis and cardiolipin remodeling during differentiation. For each week, the value presented here in the mean of n=3 replicates.

Figure 3a shows the OPLS analysis of lipids which revealed distinct clusters of lipid samples corresponding to differentiation status (differentiated cells) along positive T-score and control cells along negative T-scores. The separation along orthogonal T-score was less apparent.

**Fig. 3.**
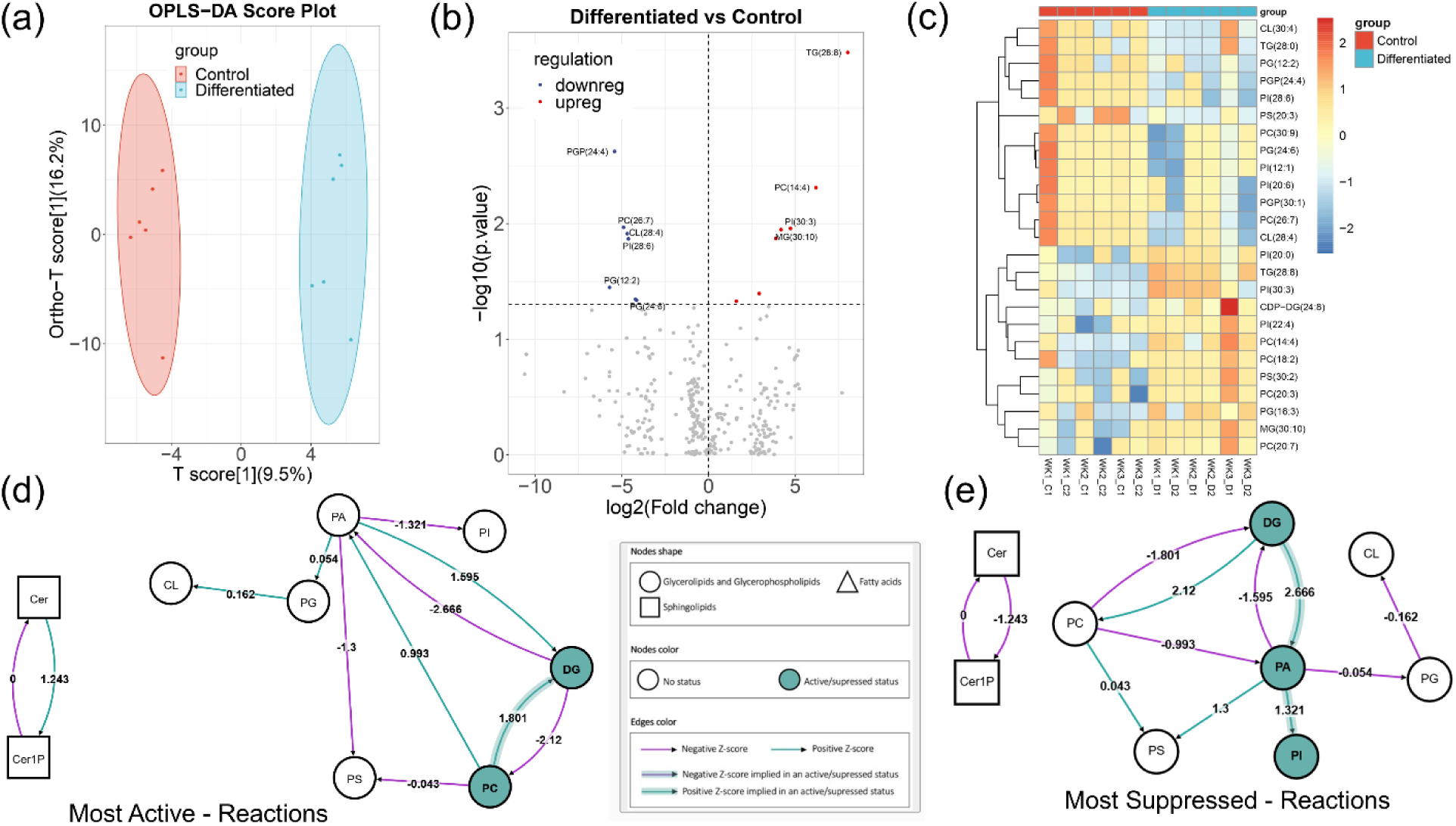
Lipidomic profiling reveals reprogramming of lipid metabolism during osteogenic differentiation: (a) OPLS-DA score plot showing distinct cluster for control and differentiated cells; (b) Volcano plot showing the significant upregulated (red) and downregulated (blue) lipid species for differentiated cells compared to stromal cells; (c) hierarchical clustering heatmap for control and differentiated stem cells. Blue color shows downregulated and red color signify upregulated lipid species; (d, e) biochemical Pathway analysis showing the formation and degradation of significant lipids with most active reactions (d) and most suppressed reactions (e). Significance was determined at P≤0.05.

Using a volcano plot (Fig. 3b), we identified differentially expressed upregulated lipid species (red color) and downregulated (blue color) lipid species for differentiated stem cells vs control (*p* ≤ 0.05, *FC* = 1). We identified top upregulated lipid species such as PGP, PC, CL, PI and PG and top downregulated lipid species such as TG, PC, PI, MG in the differentiated stem cells. The clustered heatmap analysis (Fig. 3) shows the upregulated lipids in red and the downregulated ones in blue.

The pathway analysis (Fig. 3d and 3e) of the lipids was performed using BioPAN. The active reaction chains (Fig. 3d) of PC→ DG and PA → DG, with respective Z-scores of 1.801 and 1.595. These reactions were catalyzed by genes such as PLPP1, PLPP2, and PLPP3, suggesting a robust activation of DG in the differentiated cells. The upregulation of these reactions indicates a significant shift towards lipid signaling pathways that are essential for the maintenance and functionality of differentiated cells. In addition, the reaction chain Ceramide (Cer) → Ceramide-1-phosphate (Cer1P) with Z-score of −1.243, was activated, indicating a potential downregulation of ceramide signaling pathways in differentiated cells. The reaction chain of DG→PA→PI with Z-score 3.987 and the reaction chain of DG→PA→PS with Z-score 3.966 (Fig. 1e) shown be to be suppressed in differentiated cells. These reaction chains show the significant downregulation of phosphoinositide and glycerophospholipid metabolism in differentiated cells.

To pinpoint how membrane lipids are rewired during osteogenic differentiation, we profiled the phospholipid classes and integrated enzyme expression with orthogonal LC–MS and Raman readouts. Heatmaps revealed clear separation of groups for (Fig 4(a-d)) PS, PC, PI and TG. In PS, the PUFA fraction increases by ∼15–20%, with MUFA decreasing ∼10–15% and SFA decreasing by ∼50%. PC shows a decrease by ∼50-60% in SFA and overall decrease as well. PI exhibits an increase of SFA by ∼40% while both PUFA and MUFA decease by ∼10-15% and overall decrease. In contrast, TG declines sharply in abundance (∼30-40% down) of SFA, indicating reduced routing into storage. Enzyme patterns align with these fluxes: PISD is upregulated (supporting PS→PE conversion), CDIPT is upregulated (feeding the PI branch), while PEMT is downregulated (limiting PE→PC); concurrently, DGAT2 are upregulated and DGAT1 are downregulated. Orthogonal lipid scores corroborate these trends: LC–MS (Fig. 4(k-n)) shows decreases in PS, PC and PI and increase in TG in differentiated groups from week 1 to week 3. Raman DCLS (o–r) reproduces the same directionality for PS, PC.

**Fig. 4.**
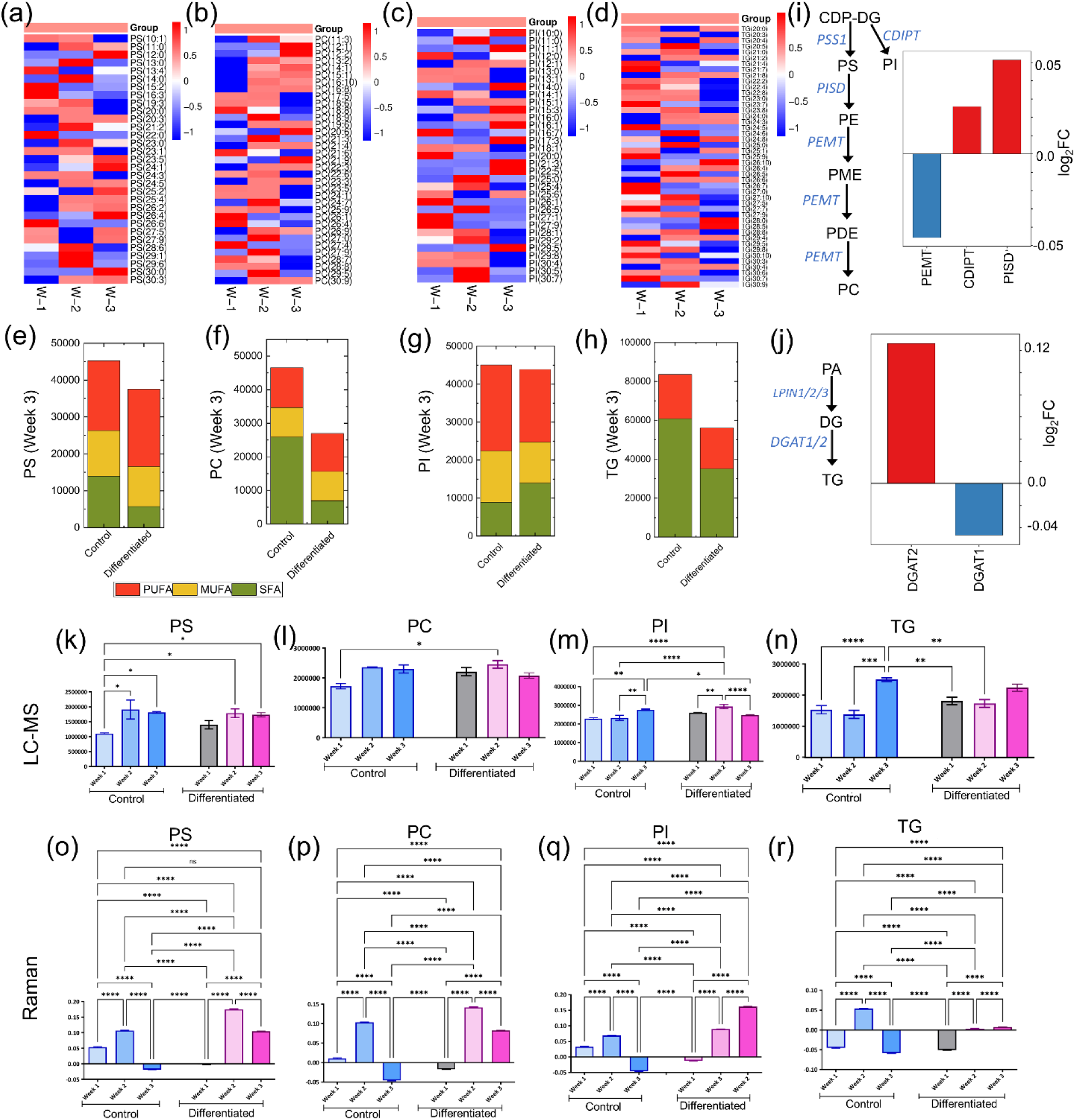
Class-specific remodeling of phospholipids during osteogenic differentiation: (a–d) heatmaps of individual species for (a) PS, (b) PC, (c) PI, and (d) TG using the FC (Diff. vs. Control) mean values at week 1, 2 and 3. The mean values are calculated based on n = 3 replicates at each week; (e–h) corresponding stacked-bar summaries of each class, depicting the fractional contribution of PUFA, MUFA, and SFA in control vs. differentiated groups in week 3 (maturation phase); (i) differential expression of enzymes governing canonical conversion of CDP to CDP-DG→PI using Kennedy pathway. The expression of PEMT, CDIPT, PISD found in our RNAseq data andtheir corresponding log_2_FC (Diff. vs. Control). Positive (red) corresponds to upregulation and negative log_2_FC value (blue) corresponds to downregulation of the transcriptome in the differentiated group; (j) The transcriptomic log_2_FC expression of neutral-lipid flux regulators (DGAT1/2) in the differentiated cells compared to the control cells. Time-dependent lipid score comparison during osteogenic differentiation for (k) PS, (l) PC, (m) PI and (n) TG using LC-MS (k-n) and the corresponding values measured using Raman DCLS analysis (o-r).

The Gene Ontology enrichment analysis (Fig. 5a) and network visualization (Fig. 5b) were performed to understand the interplay between structural remodeling, metabolic shifts, and signaling pathways highlights the complexity of osteogenesis. The dot plot represents Gene Ontology (GO) enrichment for differentially expressed genes, where the x-axis indicates the enrichment score (-log10 of the p-value), and the y-axis lists biological processes. Dot size corresponds to the number of associated genes, while color intensity represents statistical significance, with red indicating highly enriched pathways. Among the top enriched categories were bone remodeling, extracellular matrix (ECM) organization, and extracellular structure organization, highlighting the structural transformations required for osteoblast maturation. These processes are essential for collagen deposition, mineralization, and overall skeletal integrity. Interestingly, the enrichment of bone resorption—typically linked to osteoclast activity—suggests a coordinated signaling mechanism between osteoblasts and osteoclasts during bone turnover. Additionally, the enrichment of cell chemotaxis points to increased migratory behavior, which may aid in the spatial positioning of differentiating cells and the organization of newly forming bone tissue.

**Fig. 5.**
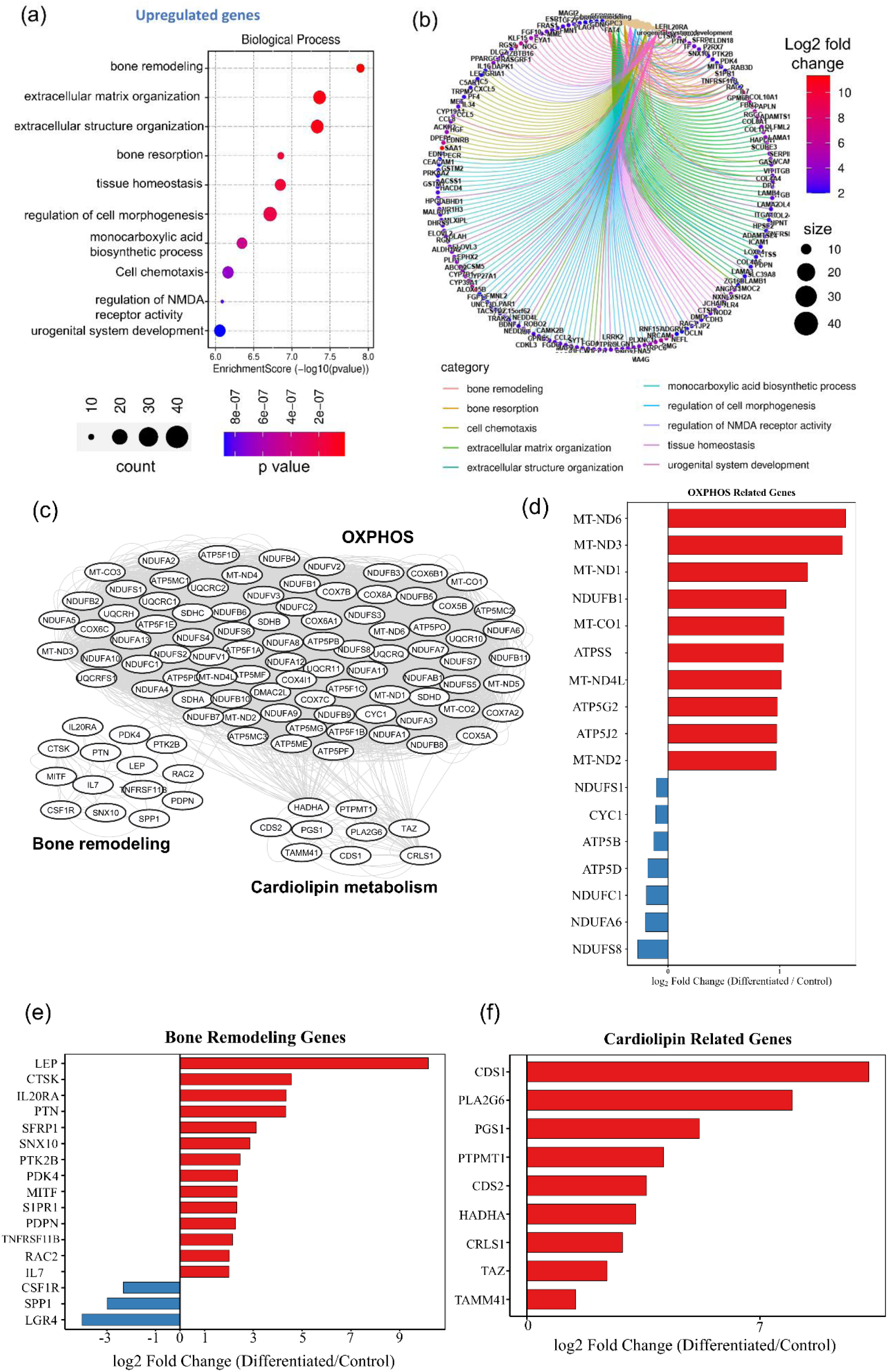
Enrichment and network analysis of biological processes in osteogenic differentiation: (a) Gene Ontology (GO) enrichment analysis of biological processes using bubble plots showing the top 10 terms for differentiated cells. The dot size (count) represents the number of genes that are within the specific GO term. The color shows p-value; (b) GO enrichment network visualization shows the intersecting genes that are enriched in bone remodeling and ECM organization; (c) STRING PPI network of the genes encoding oxidative phosphorylation (OXPHOS) (beige) with bone-remodeling markers (green) and cardiolipin-metabolism genes (pink). (d–f) Bar charts of log2 fold change (Differentiated/Control) for representative genes found in our RNAseq data that were used for the PPI network in (c): (d) OXPHOS genes (mostly upregulated), (e) bone-remodeling genes (mixed regulation), and (f) cardiolipin-pathway genes (upregulated).

Gene Ontology (GO) enrichment analysis revealed that several biological processes are significantly involved in osteogenic differentiation. Among these, metabolic pathways, particularly those related to monocarboxylic acid biosynthesis, were notably enriched. This aligns with our Raman spectroscopy (Fig. 7) and lipidomics data (Fig. 1 and 3), which show increased levels of mitochondrial lipids such as cardiolipin and. These results suggest that changes in lipid metabolism not only support the energy needs of differentiating cells but may also play a regulatory role in osteoblast function. Other enriched pathways, including tissue homeostasis and cell morphogenesis, highlight the close relationship between metabolic activity and structural changes during differentiation. Less prominent terms, such as regulation of NMDA receptor activity and urogenital system development, may reflect background gene activity or secondary roles of stem cell-related genes.

The network visualization further illustrates how these biological processes are connected through differentially expressed genes. Genes involved in bone remodeling and extracellular matrix organization show strong interconnectivity, emphasizing their importance in osteogenesis. The circular layout places biological processes at the center, with genes arranged around them. Node size indicates the number of genes linked to each process, and edge colors represent functional categories, providing a clear overview of the molecular interactions driving osteogenic differentiation.

To understand bone-remodeling, oxidative phosphorylation (OXPHOS) and cardiolipin metabolism, we used a STRING protein-protein interaction (PPI) analysis (Fig. 5c). This analysis grouped the genes that encode enzymes into distinct clusters that were consistent with the known roles of the proteins in biological process of the cells. Proteins involved in OXPHOS (Fig. 5d) in our RNAseq data were clustered together and showed a high degree of connectivity with bone remodeling (Fig. 5e) and cardiolipin metabolism (Fig. 5f). For bone remodeling, the highly connected genes were mostly upregulated (LEP, CTSK, IL20RA, PTN, SFRP1, SNX10, PTK2B, PDK4, MITF, S1PR1, PDPN, TNFRSF11B, RAC2, IL7) and few downregulated (CSF1R, SPP1 and LGR4). For Cardiolipin metabolism, interconnected genes with other two clusters are CDS1, PLA2G6, PGS1, PTPMT1, CDS2, HADHA, CRLS1, TAZ and TAMM41 which are all upregulated. The results match with stacked bar plots of fatty acid chains, heat map and circos plots for cardiolipin.

Among these enzymes, disruption of TAMM41 (mitochondrial CDP-DAG synthase), PGS1 and PTPMT1 (PGP synthesis/dephosphorylation), and CRLS1 (CL synthase) lowers CL and compromises respiratory-chain organization and OXPHOS [30–32]. Mature CL species are then established and maintained by TAZ/tafazzin remodeling, HADHA (MLCL acyltransferase) and PLA2G6/iPLA2β (repair of damaged CL), linking acyl-composition control to mitochondrial performance and disease (e.g., Barth syndrome)[33–35]. Finally, CDS1/2 at the ER supply CDP-DAG for PI and broader phospholipid flux that interfaces with mitochondrial CL synthesis, coordinating membrane biogenesis with cellular metabolism [36].

Fig. 6 integrates gene expression, protein localization, matrix mineralization, and metabolic imaging to track differentiation from week 1 to week 3. By qPCR (Fig. 6f), early markers ALPL and BMP4 rise strongly, RUNX2 shows a modest but significant increase, and late marker Osteocalcin surges by week 3 (on the order of 10²–10³), whereas BMP7 and CALCR remain unchanged indicating progression toward mature osteoblasts. These transcriptional trends are mirrored functionally: Alizarin Red S staining reveals sparse calcium deposits at week 1 but extensive mineral fields by week 3 (Fig. 6a), with integrated intensities increasing ∼2x over matched controls and many significant comparisons (p < 0.001; Fig. 6b). Concurrently, immunofluorescence shows pronounced F-actin (Phalloidin) structures and nuclear (Hoechst) organization and perinuclear Osteocalcin localization in differentiated cells (Fig. 6c), and quantification confirms a ∼10% rise in Osteocalcin signal relative to controls (Fig. 6d). Hyperspectral imaging captures spectrally distinct, heterogeneous microdomains in differentiated samples (Fig. 6e), consistent with lipid redistribution and increasing mineral complexity.

**Fig. 6.**
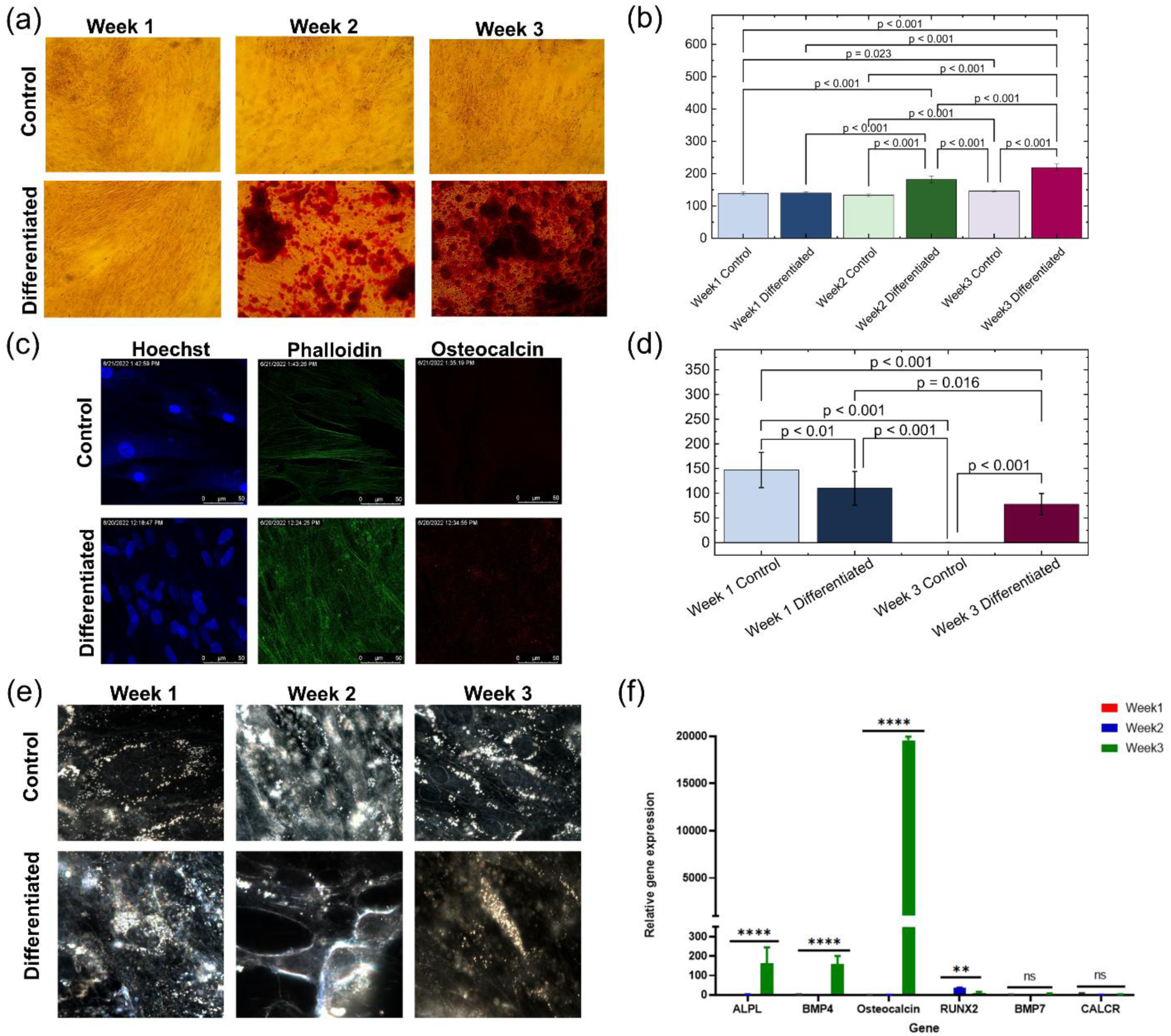
Evidence of osteogenic differentiation: (a) alizarin Red staining images showing increased calcium deposition in differentiated cells compared to control clearly seen in week 2 and week 3 of culture; (b) intensity comparison of alizarin red staining at different weeks of culture for control and differentiated samples; (c) representative fluorescence images of cells stained with Hoechst (nuclear), Phalloidin (F-actin), and Osteocalcin (osteogenic marker) at week 3 of culture; (d) intensity comparison of Osteocalcin at week 1 and week 3 of culture for control and differentiated samples; (e) hyperspectral imaging (HSI) shows spatial and spectral variations in osteogenic microenvironments and (f) quantitative PCR analysis of relative osteogenic gene expression (ALPL, BMP4, Osteocalcin, RUNX2, BMP7, and CALCR) at weeks 1, 2, and 3 for differentiated versus control stem cells.

We further assessed osteogenic differentiation using label-free dark-field microscopy coupled with hyperspectral imaging (HSI). The spectral library plot (Fig. 7b) shows reflectance spectra for eight unique endmembers (EMs), each exhibiting distinct spectral signatures. To obtain HSI spectral fingerprints from differentiated stem cells, we applied the Spectral Angle Mapper (SAM) method, which identified eight discrete EMs. The percentage classification reflects the proportion of pixels (from ∼500,000 spectra in each image corresponding to one spectrum per pixel) assigned to each EM; notably, each pixel contains both spatial and spectral information. Of these, EM-5 (SAM5) accounts for ∼10% of total scattering in week 3 differentiated cells, signifying marked bone matrix deposition. In contrast, EM-2 (SAM2) and EM-3 (SAM3) are detectable in control cells at all time points, as well as in differentiated cells during weeks 1 and 2. However, EM-5 becomes prominent only in week 3 differentiated samples. A comparative analysis of EM-1 (SAM1) and EM-5 abundance between control and differentiated groups over weeks 1–3 (Fig. 7c–d) shows elevated levels of both components in differentiated cells across all weeks. Notably, EM-5 is significantly more abundant in week 3 differentiated samples compared to week 3 controls, consistent with increased bone matrix formation.

**Fig. 7.**
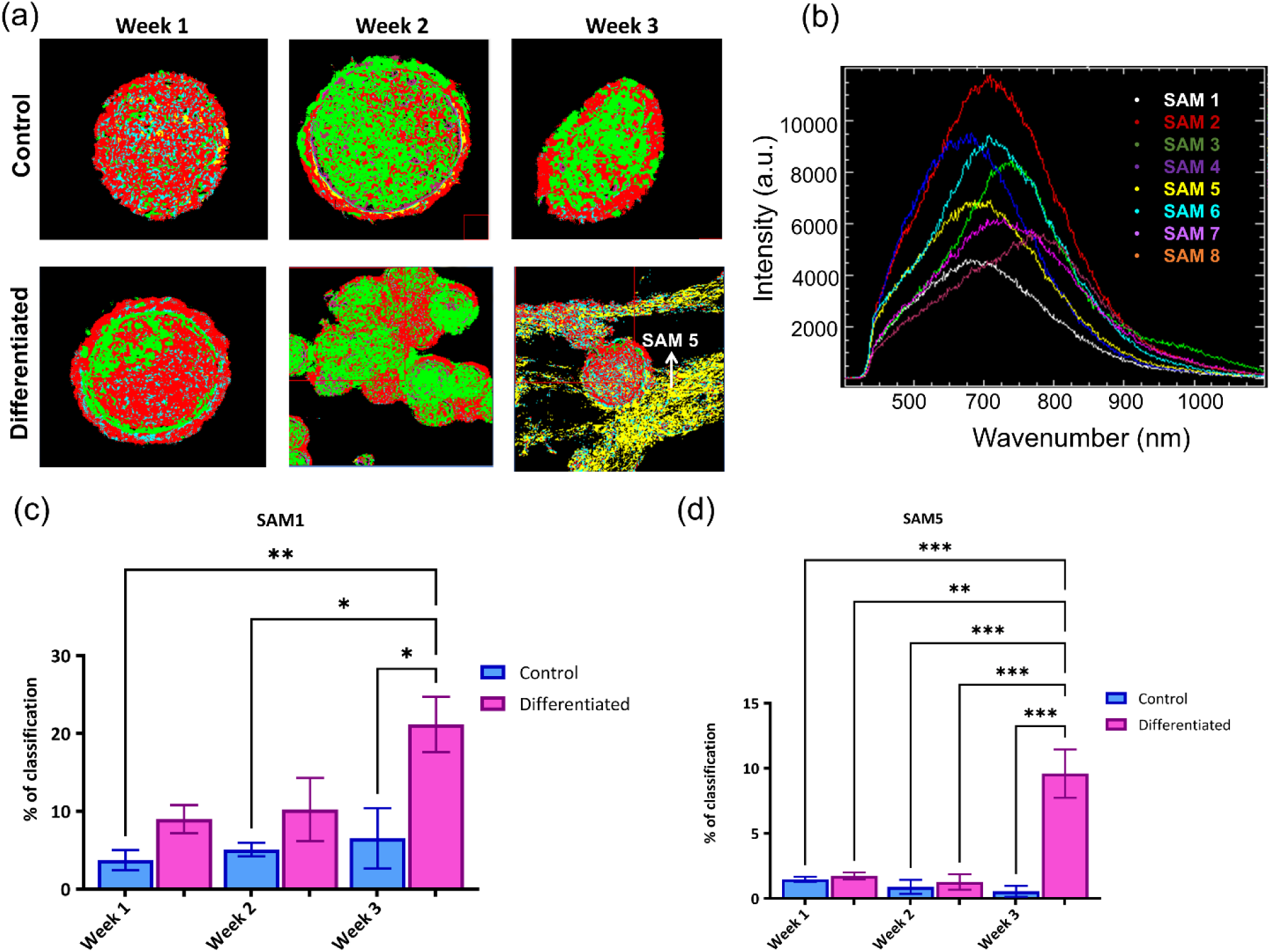
Hyperspectral imaging (HSI) analysis revealing spatial and spectral chemical distribution during osteogenic differentiation: (a) HSI-derived spatial classification maps of different chemical components, illustrating the heterogeneous distribution of spectral angle mapping (SAM) classes within differentiated and control cell; (b) spectral library plots showing the characteristic scattering profiles of key cell components and comparison of % of cells with (c) SAM 1 and (d) SAM 5 chemical components in control vs. differentiated cells for week 1, week 2 and week 3 of cell culture. SAM5 is associated with the calcium matrix deposition (yellow color) shown in (a) differentiated cell at week 3.

To find the chemical signature of the matrix in Fig. 7a (shown by SAM 5), we performed Raman microscopy experiments on these cells and showed mean of 10 spectra from control and differentiated cells of week 3 (Fig. 8a). To show the distribution of lipid species, we performed Raman mapping. Fig. 8 provides a detailed assessment of cardiolipin distribution and accumulation during osteogenic differentiation using a combination of Raman spectroscopy, LC-MS quantification, and DCLS-based spatial mapping. In Fig. 8e, Raman spectra show a prominent peak around 992 cm⁻¹, characteristic of cardiolipin [19], and a nearby β-tricalcium phosphate (β-TCP) peak at ∼950 cm⁻¹ [20] and both significantly elevated in differentiated cells compared to controls. These findings indicate not only increased mitochondrial lipid content but also concurrent mineralization during osteogenesis. The overlaid spectra and the inset bar graph visually confirm stronger signal intensities corresponding to cardiolipin in differentiated cells. Fig. 8c and 8d displays brightfield microscopy images aligned with their respective Raman DCLS maps across control and differentiated groups at weeks 1, 2, and 3.

**Fig. 8.**
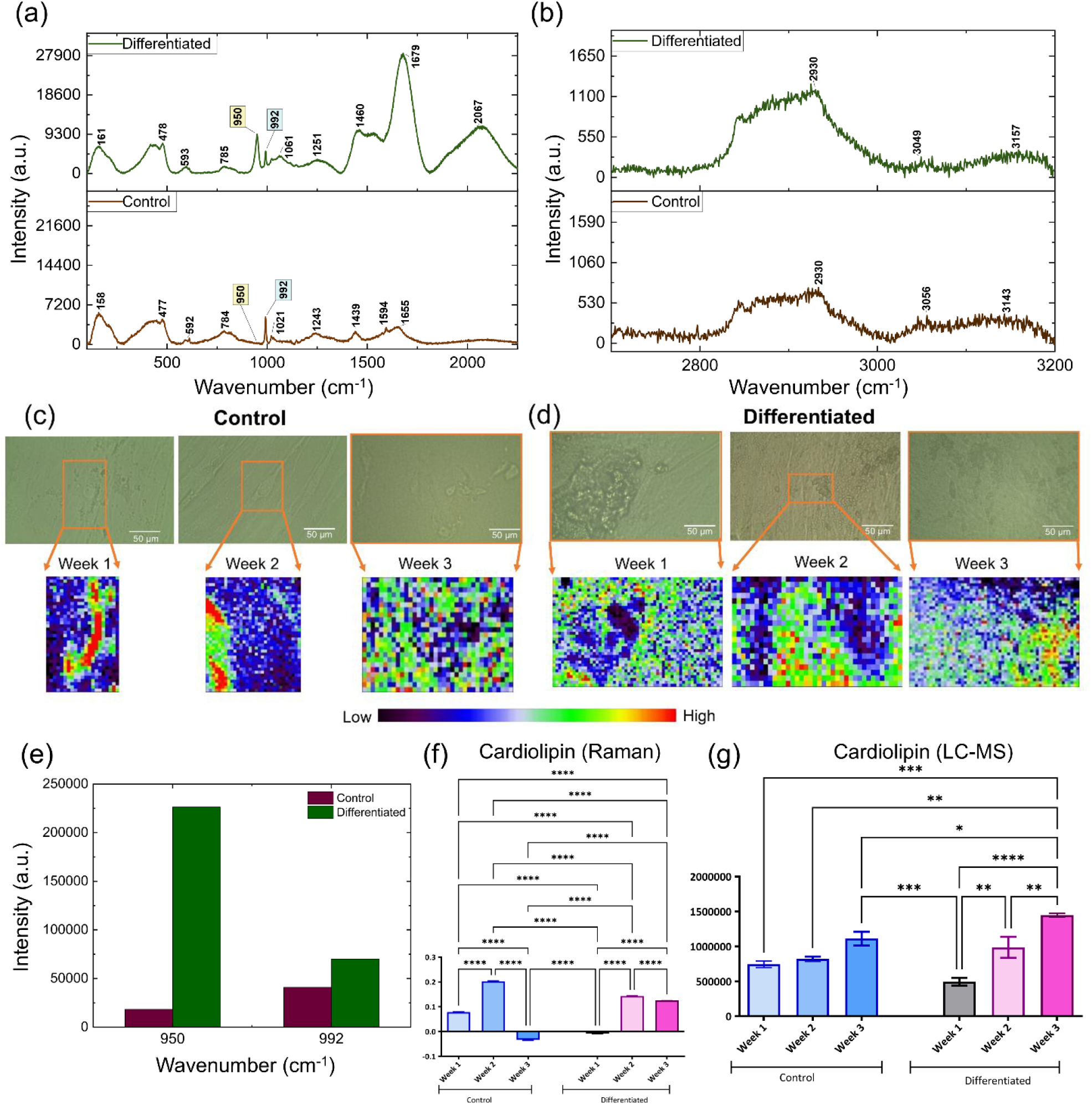
Raman analysis of cardiolipin distribution and quantification in osteogenic differentiation (week 3): (a, b) comparison of Raman spectra from control and differentiated cells in week 3 in the wavenumber range of (300 – 2200) cm^-1^ and (2600 – 3200) cm^-1^ respectively; brightfield images of Week 1, 2 and 3 samples of (c) control and (d) differentiated groups and their corresponding DCLS Raman maps of Cardiolipin; (e) Raman intensity comparison of β-tricalcium phosphate (950 cm^-1^) and Cardiolipin (992 cm^-1^). Cardiolipin quantification from (f) Raman and (g) LC-MS data for control and differentiated cells in week 1, 2 and 3 of cell culture.

Quantitative data from both Raman (Fig. 8f) and LC-MS (Fig. 8g) analyses confirm this trend: cardiolipin levels stay low in controls but rise significantly in differentiated cells from week 2 onward. The strong consistency across both techniques reinforces the idea that cardiolipin accumulation supports the increasing energy demands of osteoblast maturation and matrix formation.

Fig. 9 integrates second harmonic generation (SHG), two-photon fluorescence (TPF), and fluorescence lifetime imaging microscopy (FLIM) to characterize metabolic and structural changes during osteogenic differentiation. TPF (red) detects autofluorescent metabolic cofactors (e.g., NADH, FAD), while SHG (green) captures collagen architecture, revealing enhanced collagen deposition and autofluorescence in differentiated cells by week 3 which is evidence of matrix maturation and metabolic activation.

**Fig. 9.**
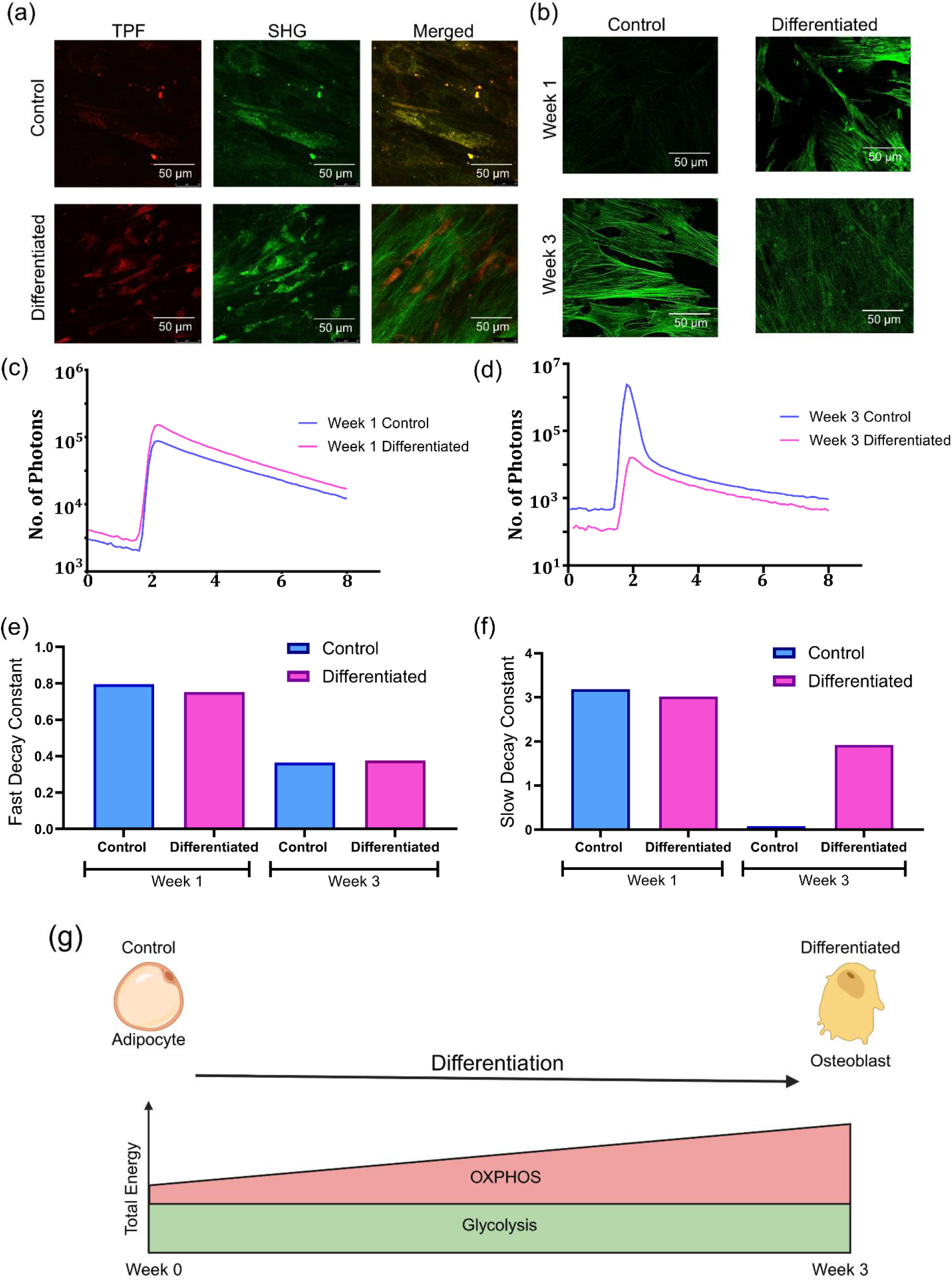
Metabolic imaging of the control and differentiated osteogenic cells: (a) representative two-photon fluorescence (TPF) and second harmonic generation (SHG) microscopy imaging of control vs. differentiated cells in week 3; (b) fluorescence lifetime imaging (FLIM) for control vs. differentiated cells for week 1 and week 3 of culture. Mean fluorescence lifetime spectra of control vs. differentiated cells in (c) week 1 and (d) week 3. Variation of fluorescence lifetime for the (e) fast and (f) slow decay constants for control and differentiated cells in week 1 and week 3; (g) Schematic representation of adipocyte transforming into osteoblast showing transition from glycolysis to OXPHOS while undergoing differentiation in week 1 to week 3.

FLIM analysis in Fig. 9(c–f) provides a deeper look into the metabolic changes associated with osteogenic differentiation by measuring the fluorescence lifetimes of endogenous fluorophores, primarily NADH. In Fig. 9b, FLIM images show clear visual differences between control and differentiated cells at both week 1 and week 3. Differentiated cells exhibit longer average lifetimes, particularly in week 3, suggesting a shift in the balance from free NADH (associated with glycolysis [39, 40]) to protein-bound NADH (indicative of oxidative phosphorylation, OXPHOS [39, 40]). FLIM maps revealed a shift of NAD(P)H toward more protein-bound states (Fig. 9b): the lifetime was slightly higher in differentiated cells at week 1 (∼10–15%) and diverged further by week 3 (∼20–40%; Fig. 9c–d). Fast and slow lifetime constants were calculated using two-phase exponential decay function with time constant parameters.

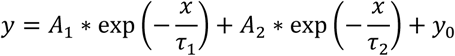

Where *y*_0_is the offset, *A*_1,2_ are amplitudes, and τ_1_and τ_2_are the fast and slow decay constants, respectively. Fig. 9(e–f) quantifies the FLIM components of NADH to track metabolic reprogramming during osteogenesis. The fast lifetime (Fig. 9e) reports free NADH, which predominates under glycolysis, whereas the slow lifetime (Fig. 9f) reflects protein-bound NADH linked to oxidative phosphorylation (OXPHOS) [37]. Week 1, both control and differentiated cells exhibit relatively high fast-lifetime values which are consistent with a glycolysis-biased state typical of proliferative, undifferentiated cells with only minor differences between groups. By Week 3, the fast decay constant did not change between control vs. differentiated cells. The slow decay constant for control vs. differentiated increased (∼2x) in week 3 signifying a shift towards OXPHOS. Together, these changes indicate a pronounced shift toward OXPHOS in differentiated cells, consistent with increased mitochondrial activity needed for extracellular-matrix production and mineralization during late-stage osteoblast maturation.

## DISCUSSION

This study demonstrates that the osteogenic differentiation of human adipose-derived stem cells is underpinned by a significant metabolic reprogramming, where the remodeling of the mitochondrial phospholipid cardiolipin (CL) acts as a central regulatory event. Our integrated, multi-modal analysis reveals that a coordinated upregulation of the cardiolipin biosynthetic and remodeling pathways drives a shift from glycolysis to oxidative phosphorylation (OXPHOS) [29, 38]. This bioenergetic enhancement, in turn, fuels the high energy demands of extracellular matrix (ECM) synthesis and mineralization [39]. This transition is consistent with previous findings showing that progenitor cells initially rely on glycolysis for rapid ATP generation to support proliferation, whereas mature osteoblasts increase their dependence on mitochondrial respiration to sustain collagen production and mineral deposition [4, 40]. Furthermore, alterations in mitochondrial cholesterol levels have recently been implicated in modulating OXPHOS efficiency, underscoring the tight interplay between lipid metabolism, mitochondrial function, and osteogenic progression [13, 41].

Our lipidomic and transcriptomic data provide a detailed molecular blueprint of this process. We identified a directed metabolic flux along the PA→CDP-DG→PGP→CL biosynthetic axis, characterized by an expansion of the PA pool and a relative contraction of CDP-DG and PGP intermediates. This was accompanied by the transcriptional upregulation of key enzymes controlling CL biosynthesis (*CDS1/2*, *PGS1*, *CRLS1*) and acyl chain remodeling (*TAZ*, *HADHA*, *PLA2G6*). Critically, the CL pool in differentiated cells became significantly enriched in highly unsaturated species, a molecular signature known to optimize the assembly and stability of respiratory chain super complexes, thereby enhancing OXPHOS efficiency. This rewiring appears to be a concerted cellular strategy, as lipid precursors were diverted from neutral lipid storage pathways toward membrane biogenesis to support this mitochondrial expansion [36].

We obtained direct functional evidence of this metabolic shift using fluorescence lifetime imaging microscopy (FLIM). By measuring the autofluorescence decay of NAD(P)H, we observed a pronounced increase in the slow lifetime component (τ_2_) in differentiated cells, particularly by week 3. This signifies a larger fraction of protein-bound NAD(P)H, a hallmark of increased OXPHOS activity relative to glycolysis [42, 43]. This finding provides a crucial link, demonstrating that the observed molecular remodeling of CL translates directly into a tangible enhancement of mitochondrial respiration at the cellular level. Raman spectroscopy and LC-MS further corroborated these findings, showing a significant accumulation of cardiolipin that spatially correlated with areas of active differentiation and mineralization.

The bioenergetic gain from enhanced OXPHOS is directly coupled with the functional outputs of osteogenesis. The timing of the metabolic shift, confirmed by FLIM, coincides with the increased expression of late-stage osteogenic markers like Osteocalcin and the extensive deposition of calcium phosphate minerals and organized collagen, as shown by qPCR, Alizarin Red staining [44], and SHG microscopy [45]. The gene ontology and network analyses further solidify this link, revealing a tightly interconnected regulatory module comprising OXPHOS, CL metabolism, and bone remodeling genes [29, 46].

In conclusion, our work integrates lipidomic profiling with functional metabolic and biochemical imaging to construct a comprehensive model of osteogenesis. Our results demonstrate that cardiolipin remodeling is a critical checkpoint that synchronizes the cell’s energy production with the demands of extracellular matrix production. Modulation of CL biosynthetic or remodeling enzymes could offer a novel strategy for enhancing bone regeneration in clinical contexts.

## MATERIALS AND METHODS

### hASCs Isolation and Osteogenic Differentiation

All materials required for osteogenic cell culture were sourced from commercial suppliers (primarily Sigma-Aldrich, St. Louis, MO, and Fisher Scientific, Hampton, NH). Experiments involving human tissue adhered to protocols approved by the Western Institutional Review Board and the Pennington Biomedical Research Center (PBRC) ethics committee. The experimental protocol was reviewed and approved by Louisiana State University’s Institutional Biosafety Committee (IBC) with approval # 24007. Human adipose-derived stem cells (hASCs) were obtained from de-identified lipoaspirate samples donated with informed consent. Within a day after collection, the adipose material was washed repeatedly (four cycles) in pre-warmed phosphate-buffered saline (PBS, 37 °C) to remove circulating blood components. The methodology used for the separation and expansion of hASCs has been previously reported in detail [47, 48]. After enzymatic digestion, the cell suspension was centrifuged at 300g for 5 min at room temperature (≈25 °C) to remove remaining collagenase. The resulting cell pellet was transferred into stromal growth medium and expanded under standard culture conditions (37 °C, 5% CO₂, humidified atmosphere). To initiate osteogenic differentiation, cultures were switched to osteoinductive media (OSTEOQUAL, LaCell LLC, New Orleans, LA) and maintained for 21 days, with media refreshed approximately every three days. Control cultures were grown in stromal medium composed of DMEM supplemented with 10% fetal bovine serum (FBS) and 1% penicillin–streptomycin.

### Raman Imaging Osteogenic Differentiated Cells

Raman spectral data and imaging of hASCs were obtained using a Renishaw inVia Reflex Raman Microspectroscope (Renishaw, UK) equipped with a 50× long working distance objective. Spectra were collected with a 785 nm excitation laser, an acquisition time of 20 s, and a spectral window spanning 200–3200 cm⁻¹. Raman mapping was performed in Map Image Acquisition mode, while Raman imaging utilized a 1200 l/mm grating (633/780 nm), with a 2 s integration time per spectrum in static mode, centered at 1250 cm⁻¹ and spanning a wavenumber range of 665–1773 cm⁻¹. Calibration of all spectra was performed against the standard silicon reference peak at 520 cm⁻¹. Comparative Raman analyses were conducted on both control and osteogenically induced cells at weeks 1, 2, and 3 of culture. Raman images were further processed using the direct classical least squares (DCLS) algorithm [49], which enables quantification of molecular changes when reference spectra of specific components are available. In this study, lipid reference spectra were used to monitor remodeling of lipid profiles on the cell surface before and after osteogenic differentiation.

### Multiphoton Microscopy of Osteogenic Stem Cells

Two-photon fluorescence (TPF) and second harmonic generation (SHG) imaging of stem cells undergoing osteogenic differentiation were conducted on a Leica SP8 resonant-scanning multiphoton confocal platform (Leica Microsystems). The system was powered by a Spectra Physics Mai-Tai tunable femtosecond near-infrared laser source (690–1040 nm), and imaging was performed using a 63× oil-immersion objective. Endogenous metabolic fluorophores, specifically NADH and FAD, were visualized using a Ti:Sa femtosecond laser that operated at a repetition frequency of 4000 MHz with ∼70 fs pulse width. NADH signals were excited at 750 nm, and emission was recorded within the 320–430 nm spectral window. For FAD, excitation was set to 860 nm, and fluorescence was detected using a 486–506 nm bandpass filter.

### Alizarin Red Staining

hASCs, cultured with or without osteogenic induction media for 7, 14, and 21 days, were rinsed with PBS and fixed in 4% paraformaldehyde for 30 minutes at room temperature. Following fixation, the cells were washed again with PBS and treated with 60% isopropanol for 5 minutes. The isopropanol was aspirated, and cells were incubated with Alizarin Red solution for 15 minutes to assess mineral deposition. After staining, cells were rinsed 2–3 times with PBS, and optical images were acquired at 10x magnification. Image analysis was performed using ImageJ software, where images were thresholded by color and hue, and quantified based on area and intensity within defined regions of interest.

### Fluorescence Imaging

For fluorescence imaging of osteogenic stem cells, Hoechst (blue) (λ_ex_: 460/λ_em_: 490 nm) stained nuclei by binding to A-T regions of DNA, while Phalloidin (green) (λ_ex_: 488/λ_em_: 510 nm) labeled the cytoskeleton. Runx2 (red), an early osteogenesis marker, was highly expressed in immature osteoblasts (Week 1) and downregulated in mature osteoblasts (Week 2), localizing around the nucleus. Osteocalcin (red spots), a bone-forming protein, was used to identify mature osteoblasts through immunostaining. These markers collectively provided insights into osteoblast differentiation and commitment. Imaging was conducted using a Leica TCS SP8 confocal fluorescence microscope (63x/1.20 NA water immersion), and data were analyzed with ImageJ.

### Lipidomics data analysis

LINT-web (http://www.lintwebomics.info/single) was used for the analysis of lipidomics data [51]. The lipid expression data in .csv format was uploaded to the LINT-web tool. Under the data preprocessing step, lipid species with more than 70% missing values were removed before the analysis. In order to remove the variance between the samples we utilized the median norm along with the log transformation and autoscaling. The Orthogonal partial least squares discriminant analysis (OPLS-DA) and principal component analysis (PCA) were studied to perform the dimensionality reduction on the samples. The differential expression of the lipids was studied using student t-test. The results were exported in the various forms of the graphs like cumulative graph of the lipids, scores plot, stacked histogram, heatmap and volcano plot of lipid differences between the control and differentiated samples. The lipid ontology analysis was carried out by utilizing Lipid Ontology (http://lipidontology.com/) to classify the most highly enriched lipid ontology terms among all the differentially expressed lipids.

### Biopan Pathway analysis

The lipid expression data in .csv format was submitted to the BioPAN pathway analysis tool, which was designed specifically for the lipidomics data (https://www.lipidmaps.org/biopan/) [52]. BioPAN determines the statistical scores for all the potential lipid pathways and their reactions, which can be active or suppressed in differentiated samples compared to the control samples. In brief, the workflow of the BioPAN uses the Z-score as a statistical measure that considers both the standard deviation and the means to estimate the normal distribution of the lipidomics data and find out the pathway or a lipid reaction that can be changed at a significant p-value, i.e., less than 0.05. The detailed calculation of the Z-score was reported by Gaud et al. [53].

### Hyperspectral Imaging

Dark-field hyperspectral imaging (HSI) was employed to study osteogenic differentiation of human adipose-derived stem cells (hASCs) in a non-invasive manner. High-resolution spectral data were collected using a CytoViva Hyperspectral Microscope operating in the visible–near-infrared range (400–1000 nm). Illumination in dark-field mode was achieved by directing light from a high-intensity halogen source through a liquid light guide at oblique angles, providing efficient scattering contrast. Cells were cultured on glass slides, covered with coverslips, and imaged using a 100× oil immersion objective, which enabled precise visualization of lipid droplet formation.

During acquisition, more than 500,000 scattering spectra per image were recorded and processed using the Spectral Angle Mapper (SAM) machine learning approach. This method identified eight distinct spectral endmembers corresponding to ECM, bone matrix, cytoplasm, and other cellular components. The system provided a spectral resolution of 2 nm with a spatial sampling width of 25 nm per pixel. Each pixel was exposed for 0.30 seconds, and detector intensity was adjusted within a range of 1000–10,000 counts to minimize background noise and avoid saturation.

Raw spectral data were preprocessed by background subtraction and normalization, followed by Principal Component Analysis (PCA) to validate the distinctiveness of the spectral signatures. For classification, the SAM algorithm compared the spectral vector of each pixel from control and differentiated cells against reference endmember spectra. Because SAM is insensitive to brightness variations from the illumination source, radiance values were converted to absorbance and dark values were excluded prior to analysis. Pixel classification outcomes were normalized to the proportion of biomatrix present in the field of view (FOV). Importantly, the SAM approach demonstrated robustness to variations in albedo and lamp intensity, ensuring consistency and reliability in the spectral analysis.

### Gene Ontology enrichment analysis and Network visualization

The SRplot online platform (https://www.bioinformatics.com.cn/en) was used to perform Gene Ontology (GO) enrichment and network visualization analyses under GO, pathway Enrichment Analysis [54]. The RNA sequencing data-derived differentially expressed genes (DEGs) were uploaded as official gene symbols along with respective fold change (FC). The analysis was performed using a total no. of DEGs with the log2FC ≥ 2 and log2FC ≤-2. The network visualizations depicted GO term relationships with their associated genes through node size indicating linked gene numbers and edge color representing functional categories. These analyses were used to identify the major biological pathways and gene interactions that contribute to osteogenic differentiation.

### Interaction Map; CIRCOS Plot

We utilized biological processes, such as extracellular matrix organization and bone remodeling, identified through Gene Ontology (GO) analysis, to construct the protein-protein interaction map. We constructed a protein–protein interaction (PPI) network to examine the interactions between ECM-related and bone-remodeling-related genes. We screened the STRING database (v11.5) [55] using a confidence score criterion of ≥ 0.7 to make sure that the associations were strong. The lipid expression data in .csv format were preprocessed, and the circos plot of lipidomic changes during week 1, week 2, and week 3 was generated using the circlize R package [56].

### RNA Isolation qPCR

RNA was isolated from stem cell cultures using the PureLink RNA Mini Kit (Thermo Fisher Scientific, MA, USA),and yield as well as purity were assessed spectrophotometrically with a Nanodrop instrument. First-strand complementary DNA (cDNA) was generated from the extracted RNA using the High-Capacity cDNA Reverse Transcription Kit (Thermo Fisher Scientific, MA, USA) according to the manufacturer’s instructions. Quantitative PCR (qPCR) analysis was performed on an ABI 7900 real-time system with SYBR Green PCR Master Mix (Thermo Fisher Scientific, MA, USA). Primer sequences for osteogenic differentiation markers and for the internal control gene, cyclophilin B, were adapted from previously published reports. Relative expression was determined by normalizing target genes to cyclophilin B and calculating fold changes with the 2–ΔΔCT method.

### Liquid chromatography - mass spectrometry (LC-MS)

Cell pellets were disrupted by adding 250 μL of LC–MS–grade methanol and vortexing the suspension for 1 minute. The mixtures were then placed on an orbital shaker at 500 rpm for 30 minutes at room temperature. After this step, 250 μL of LC–MS–grade water was introduced, and samples were centrifuged at 12,000 rpm for 10 minutes to remove cellular debris. Supernatants were carefully collected into new microcentrifuge tubes. To extract lipids, 500 μL of HPLC–grade chloroform was added to each sample, followed by 1 minute of vortexing and another 30 minutes of incubation under the same shaking conditions. The organic phase was isolated, transferred to fresh tubes, and evaporated under a gentle stream of nitrogen. Dried extracts were then resuspended in 40 μL of 30% acetonitrile containing 0.1% formic acid.

Lipidomic profiling was carried out using an Agilent 1260 Infinity II quaternary liquid chromatography system coupled with an Agilent 6230 electrospray ionization time-of-flight (ESI-TOF) mass spectrometer (Agilent Technologies, Santa Clara, CA, USA). Data acquisition was performed in positive ion mode with a capillary voltage of 4000 V. Nitrogen was employed as the drying gas at 10 L/min and 325 °C. The fragmentor voltage was set at 150 V, and the scan range was 100–3000 m/z. Separation was achieved on an Agilent Poroshell 120 EC-C18 column (2.7 mm ID × 150 mm length, 2.7 μm particle size, end-capped). Mobile phase A consisted of water with 0.1% formic acid, while mobile phase B was acetonitrile. The flow rate was maintained at 400 μL/min. The gradient profile was programmed as follows: 0–5 min, 5% B; 5–30 min, linear ramp to 90% B; 30–35 min, hold at 90% B; 35–45 min, return to 5% B. Each injection used 30 μL of sample. Raw LC–MS data were converted to the mzData file format using MassHunter Workstation Qualitative Analysis Navigator software (version B.08.00, build 8.0.8208.0). Further processing, feature extraction, and lipid identification were conducted with MZMine 2 and MetaboAnalyst 5.0.

## ACKNOWLEDGMENTS FUNDING

This work was supported by the National Institute of General Medical Sciences of the National Institutes of Health under award number R35GM150564. We thank Nishir Mehta, Elnaz Sheikh, Sukkum Chang, and David Burk for the data analysis.

## DATA AVAILABILITY

All data used to support the findings in the paper are available from the corresponding authors upon reasonable request.

## SUPPLEMENTARY MATERIALS

None.

## REFERENCES

1. Infante, A. and C.I. Rodríguez, Osteogenesis and aging: lessons from mesenchymal stem cells. Stem Cell Research & Therapy, 2018. 9(1): p. 244.

2. Amarasekara, D.S., S. Kim, and J. Rho, Regulation of Osteoblast Differentiation by Cytokine Networks. International Journal of Molecular Sciences, 2021. 22(6): p. 2851.

3. Shen, L., G. Hu, and C.M. Karner, Bioenergetic Metabolism In Osteoblast Differentiation. Curr Osteoporos Rep, 2022. 20(1): p. 53–64.

4. Smith, C.O. and R.A. Eliseev, Energy Metabolism During Osteogenic Differentiation: The Role of Akt. Stem Cells and Development, 2020. 30(3): p. 149–162.

5. Zuk, P.A., et al., Human adipose tissue is a source of multipotent stem cells. Mol Biol Cell, 2002. 13(12): p. 4279–95.

6. Gimble, J.M., A.J. Katz, and B.A. Bunnell, Adipose-Derived Stem Cells for Regenerative Medicine. Circulation Research, 2007. 100(9): p. 1249–1260.

7. Pittenger, M.F., et al., Mesenchymal stem cell perspective: cell biology to clinical progress. npj Regenerative Medicine, 2019. 4(1): p. 22.

8. Liao, H.T. and C.T. Chen, Osteogenic potential: Comparison between bone marrow and adipose-derived mesenchymal stem cells. World J Stem Cells, 2014. 6(3): p. 288–95.

9. Komori, T., Molecular Mechanism of Runx2-Dependent Bone Development. Mol Cells, 2020. 43(2): p. 168–175.

10. Long, F., Building strong bones: molecular regulation of the osteoblast lineage. Nature Reviews Molecular Cell Biology, 2012. 13(1): p. 27–38.

11. Decker, S.T. and K. Funai, Mitochondrial membrane lipids in the regulation of bioenergetic flux. Cell Metab, 2024. 36(9): p. 1963–1978.

12. Solsona-Vilarrasa, E., et al., Cholesterol enrichment in liver mitochondria impairs oxidative phosphorylation and disrupts the assembly of respiratory supercomplexes. Redox Biol, 2019. 24: p. 101214.

13. van Gastel, N. and G. Carmeliet, Metabolic regulation of skeletal cell fate and function in physiology and disease. Nature Metabolism, 2021. 3(1): p. 11–20.

14. Lecka-Czernik, B., Marrow fat metabolism is linked to the systemic energy metabolism. Bone, 2012. 50(2): p. 534–9.

15. Zhu, S., et al., Cell signaling and transcriptional regulation of osteoblast lineage commitment, differentiation, bone formation, and homeostasis. Cell Discovery, 2024. 10(1): p. 71.

16. Akhmetshina, A., D. Kratky, and E. Rendina-Ruedy, Influence of Cholesterol on the Regulation of Osteoblast Function. Metabolites, 2023. 13(4).

17. Choi, I.A., et al., Bone metabolism – an underappreciated player. npj Metabolic Health and Disease, 2024. 2(1): p. 12.

18. Houschyar, K.S., et al., Wnt Pathway in Bone Repair and Regeneration - What Do We Know So Far. Front Cell Dev Biol, 2018. 6: p. 170.

19. Mathiowetz, A.J. and J.A. Olzmann, Lipid droplets and cellular lipid flux. Nature Cell Biology, 2024. 26(3): p. 331–345.

20. Cliff, T.S. and S. Dalton, Metabolic switching and cell fate decisions: implications for pluripotency, reprogramming and development. Current Opinion in Genetics & Development, 2017. 46: p. 44–49.

21. Suh, J. and Y.S. Lee, The multifaceted roles of mitochondria in osteoblasts: from energy production to mitochondrial-derived vesicle secretion. J Bone Miner Res, 2024. 39(9): p. 1205–1214.

22. Huang, X., et al., Pinpointing Fat Molecules: Advances in Coherent Raman Scattering Microscopy for Lipid Metabolism. Anal Chem, 2024. 96(20): p. 7945–7958.

23. Yan, C., et al., Mitochondrial quality control and its role in osteoporosis. Front Endocrinol (Lausanne), 2023. 14: p. 1077058.

24. Meshcheryakova, A., D. Mechtcheriakova, and P. Pietschmann, Sphingosine 1-phosphate signaling in bone remodeling: multifaceted roles and therapeutic potential. Expert Opin Ther Targets, 2017. 21(7): p. 725–737.

25. Emaus, K.J., et al., The role of cardiolipin in mitochondrial dynamics and quality control in neuronal ischemia/reperfusion injury. Cell Death & Disease, 2025. 16(1): p. 494.

26. Yoo, Y., et al., Role of cardiolipin in skeletal muscle function and its therapeutic implications. Cell Communication and Signaling, 2025. 23(1): p. 36.

27. Venkatraman, K. and I. Budin, Cardiolipin remodeling maintains the inner mitochondrial membrane in cells with saturated lipidomes. Journal of Lipid Research, 2024. 65(8).

28. Kopeć, M., et al., The role of cardiolipin and cytochrome c in mitochondrial metabolism of cancer cells determined by Raman imaging: in vitro study on the brain glioblastoma U-87 MG cell line. Analyst, 2024. 149(9): p. 2697–2708.

29. Hryc, C.F., et al., Structural insights into cardiolipin replacement by phosphatidylglycerol in a cardiolipin-lacking yeast respiratory supercomplex. Nature Communications, 2023. 14(1): p. 2783.

30. Blunsom, N.J., et al., Mitochondrial CDP-diacylglycerol synthase activity is due to the peripheral protein, TAMM41 and not due to the integral membrane protein, CDP-diacylglycerol synthase 1. Biochim Biophys Acta Mol Cell Biol Lipids, 2018. 1863(3): p. 284–298.

31. Zhang, J., et al., Mitochondrial phosphatase PTPMT1 is essential for cardiolipin biosynthesis. Cell Metab, 2011. 13(6): p. 690–700.

32. Gohil, V.M., et al., Cardiolipin biosynthesis and mitochondrial respiratory chain function are interdependent. J Biol Chem, 2004. 279(41): p. 42612–8.

33. Miklas, J.W., et al., TFPa/HADHA is required for fatty acid beta-oxidation and cardiolipin re-modeling in human cardiomyocytes. Nature Communications, 2019. 10(1): p. 4671.

34. Zhu, S., et al., Cardiolipin Remodeling Defects Impair Mitochondrial Architecture and Function in a Murine Model of Barth Syndrome Cardiomyopathy. Circ Heart Fail, 2021. 14(6): p. e008289.

35. Musthafa, T., et al., Altered Mitochondrial Bioenergetics and Calcium Kinetics in Young-Onset PLA2G6 Parkinson’s Disease iPSCs. Journal of Neurochemistry, 2025. 169(4): p. e70059.

36. Blunsom, N.J. and S. Cockcroft, CDP-Diacylglycerol Synthases (CDS): Gateway to Phosphatidylinositol and Cardiolipin Synthesis. Front Cell Dev Biol, 2020. 8: p. 63.

37. Skala, M.C., et al., In vivo multiphoton fluorescence lifetime imaging of protein-bound and free nicotinamide adenine dinucleotide in normal and precancerous epithelia. J Biomed Opt, 2007. 12(2): p. 024014.

38. Messina, M., F.M. Vaz, and S. Rahman, Mitochondrial membrane synthesis, remodelling and cellular trafficking. Journal of Inherited Metabolic Disease, 2025. 48(1): p. e12766.

39. Ponzetti, M. and N. Rucci, Osteoblast Differentiation and Signaling: Established Concepts and Emerging Topics. International Journal of Molecular Sciences, 2021. 22(13): p. 6651.

40. Shyh-Chang, N. and H.H. Ng, The metabolic programming of stem cells. Genes Dev, 2017. 31(4): p. 336–346.

41. Wang, B., et al., Lipid metabolism within the bone micro-environment is closely associated with bone metabolism in physiological and pathophysiological stages. Lipids in Health and Disease, 2022. 21(1): p. 5.

42. Niederschweiberer, M.A., et al., NADH Fluorescence Lifetime Imaging Microscopy Reveals Selective Mitochondrial Dysfunction in Neurons Overexpressing Alzheimer’s Disease–Related Proteins. Frontiers in Molecular Biosciences, 2021. **Volume** 8 **-** 2021.

43. Meleshina, A.V., et al., Two-photon FLIM of NAD(P)H and FAD in mesenchymal stem cells undergoing either osteogenic or chondrogenic differentiation. Stem Cell Research & Therapy, 2017. 8(1): p. 15.

44. Bernar, A., et al., Optimization of the Alizarin Red S Assay by Enhancing Mineralization of Osteoblasts. International Journal of Molecular Sciences, 2023. 24(1): p. 723.

45. Aghigh, A., et al., Second harmonic generation microscopy: a powerful tool for bio-imaging. Biophysical Reviews, 2023. 15(1): p. 43–70.

46. Schmidt, J.R., et al., Meta-analysis of proteomics data from osteoblasts, bone, and blood: Insights into druggable targets, active factors, and potential biomarkers for bone biomaterial design. Journal of Tissue Engineering, 2024. 15: p. 20417314241295332.

47. Shaik, S., et al., Effects of Decade Long Freezing Storage on Adipose Derived Stem Cells Functionality. Scientific Reports, 2018. 8(1): p. 8162.

48. Mehta, N., et al., Multimodal Label-Free Monitoring of Adipogenic Stem Cell Differentiation Using Endogenous Optical Biomarkers. Advanced Functional Materials, 2021. 31(43): p. 2103955.

49. Kelley, D.P., et al., Labelfree mapping and profiling of altered lipid homeostasis in the rat hippocampus after traumatic stress: Role of oxidative homeostasis. Neurobiology of Stress, 2022. 20: p. 100476.

50. Quinn, K.P., et al., Characterization of metabolic changes associated with the functional development of 3D engineered tissues by non-invasive, dynamic measurement of individual cell redox ratios. Biomaterials, 2012. 33(21): p. 5341–5348.

51. Li, F., et al., LINT-Web: A Web-Based Lipidomic Data Mining Tool Using Intra-Omic Integrative Correlation Strategy. Small Methods, 2021. 5(9): p. e2100206.

52. Sheikh, E., et al., Multimodal Imaging of Pancreatic Cancer Microenvironment in Response to an Antiglycolytic Drug. Adv Healthc Mater, 2023. 12(31): p. e2301815.

53. Gaud, C., et al., BioPAN: a web-based tool to explore mammalian lipidome metabolic pathways on LIPID MAPS. F1000Res, 2021. 10: p. 4.

54. Tang, D., et al., SRplot: A free online platform for data visualization and graphing. PLoS One, 2023. 18(11): p. e0294236.

55. Szklarczyk, D., et al., The STRING database in 2023: protein-protein association networks and functional enrichment analyses for any sequenced genome of interest. Nucleic Acids Res, 2023. 51(D1): p. D638–d646.

56. Gu, Z., et al., *circlize Implements and enhances circular visualization in R*. Bioinformatics, 2014. 30(19): p. 2811–2.

